# Trimethylamine-N-oxide affects cell type-specific pathways and networks in mouse aorta to promote atherosclerotic plaque vulnerability

**DOI:** 10.1101/2025.02.25.640205

**Authors:** Jenny Cheng, Michael Cheng, Satyesh Sinha, Ingrid Cely, Sharda Charugundla, Maggie T. Han, Guanglin Zhang, Zhiqiang Zhou, Sasha Gladkikh, In Sook Ahn, Graciel Diamante, Yuchen Wang, Zeneng Wang, Brian Bennett, Hua Cai, Hooman Allayee, Stanley Hazen, Aldons J. Lusis, Xia Yang, Diana M. Shih

## Abstract

**Background:** Trimethylamine-N-oxide (TMAO) has been significantly linked to atherosclerosis via several mechanisms, but its direct effect on the atherosclerosis-prone vasculature remains unclear. The objective of this study was to characterize the cell type-dependent and independent effects of TMAO on key vascular cell types involved in atherosclerosis progression *in vivo*.

**Methods:** We performed single cell RNA-sequencing (scRNAseq) on aortic athero-prone regions of female *Ldlr−/−* mice fed control Chow, high-cholesterol (HC), or HC+TMAO diets for three months to identify which aortic cell types, differentially expressed genes, and biological pathways are affected by TMAO. We also modeled cell-cell communications and intracellular gene regulatory networks to identify gene networks perturbed by TMAO feeding. Key genes and pathways were validated using primary human smooth muscle cells exposed to TMAO. Changes in the thickness of lesional fibrous caps in response to TMAO in female *Ldlr−/−* mice fed HC+TMAO versus HC diets were measured using transgelin immunostaining.

**Results:** Our scRNAseq analysis revealed that TMAO supplementation upregulated apoptotic gene signatures and downregulated extracellular matrix (ECM) organization and collagen formation genes in a subset of atherosclerosis-specific modulated vascular smooth muscle cells (vSMCs). We also identified “degradation of the ECM” as a top pathway for SMC-derived macrophage DEGs in response to TMAO. Network analyses support that macrophage-vSMC communication mediates ECM remodeling. Using human smooth muscle cells exposed to TMAO *in vitro*, we confirmed the direct effect of TMAO on regulating collagen and apoptotic genes. In agreement with the changes in these pathways that affect plaque stability, we observed a significant decrease in fibrous cap thickness in mice supplemented with TMAO.

**Conclusions:** Our results reveal the effects of TMAO on vSMCs to promote apoptosis and decrease ECM formation, and on macrophage-mediated ECM degradation in atherosclerotic lesions to in concert enhance plaque instability.

**Graphic Abstract:** 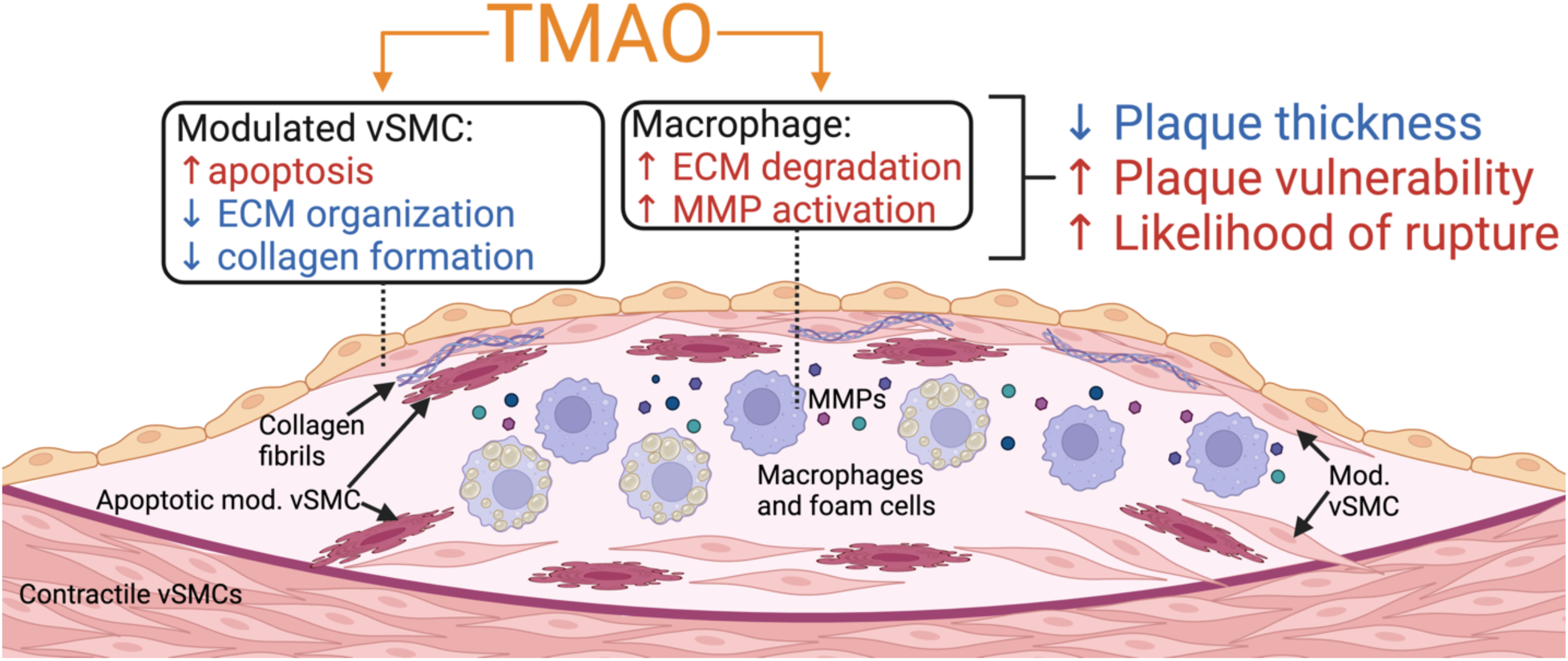

**Highlights:**

- scRNAseq of the aortic athero-prone regions in female *Ldlr−/−* mice supplemented with TMAO in the diet revealed the effect of TMAO across cell types, particularly in SMC-derived macrophages and atheroprotective modulated vSMCs.
- TMAO increases apoptotic gene signatures and reduces ECM organization and collagen formation gene signatures in modulated vSMCs *in vivo*, and *in vitro* exposure studies support a direct effect of TMAO on these genes.
- Modulated vSMC-specific gene regulatory networks enriched for apoptotic genes and ECM organization genes were organized by intracellular regulators such as Ccl19 and Tnn and extracellular regulators such as Mmp9 and Spp1 from macrophages.
- Fibrous cap thickness, a marker of atherosclerotic plaque stability, was significantly reduced in female *Ldlr−/−* mice fed HC+TMAO versus HC diets for five months.

## Introduction

The diet is a major environmental exposure that influences metabolism and shapes gut microbial composition. The gut microbiome, in turn, participates in nutrient metabolism, leading to the generation of biologically active metabolites to modulate host physiology and health^1^. Certain gut microbiota-derived metabolites have been increasingly recognized as either protective or risk factors in the development and progression of various cardiometabolic diseases. One such microbial metabolite is trimethylamine N-oxide (TMAO), which has been significantly associated with or shown to be causal for numerous cardiovascular diseases and outcomes, including atherosclerosis, abdominal aortic aneurysm, thrombosis risk, incident major adverse cardiovascular events, and heart failure^2, 3, 4, 5, 6, 7^. TMAO is derived from trimethylamine (TMA) by microbial enzyme TMA lyase initially metabolizing dietary choline, phosphatidylcholine, and L-carnitine, which are abundant in red meat and full-fat dairy products. TMA is subsequently absorbed by the host and metabolized to TMAO by hepatic flavin monooxygenase 3 (FMO3)^8^.

TMAO is an independent predictor of high atherosclerotic burden in human patients and has been demonstrated to increase atherosclerotic lesion size and foam cell formation in mice^2, 9^. Mechanistically, TMAO has been linked to atherosclerosis through several atherogenic pathways. First, TMAO alters host sterol and lipid metabolism by induction of scavenger receptors CD36 and SRA1 on macrophages^2^. This results in increased foam cell formation as well as alterations in systemic cholesterol metabolism^10, 11, 12^. Second, TMAO has also been shown to exert a proinflammatory effect on vascular cells. Aortas of *Ldlr−/−* mice fed a high-choline diet exhibited elevated inflammatory gene expression. Also, primary human aortic endothelial cells and vascular smooth muscle cells (vSMCs) treated with TMAO exhibited activation of nuclear factor-κB (NFκB) signaling and leukocytes binding to endothelial cells^13^. In endothelial cells and *ApoE−/−* mouse aortas, TMAO activated the NLRP3 inflammasome and stimulated generation of mitochondrial reactive oxygen species (ROS)^14, 15^. In vSMCs, TMAO induced nicotinamide adenine dinucleotide phosphate oxidase 4 expression and ROS production to promote vascular inflammation^16^. Third, in coronary artery disease (CAD) patients, plasma TMAO levels were significantly inversely correlated with fibrous cap thickness and positively correlated with rupture^17^. In the tandem stenosis mouse model of atherosclerotic plaque instability, plasma TMAO levels correlated with plaque instability characteristics, including inflammation, platelet activation, and intraplaque hemorrhage, whereas a reduction in TMAO enhanced carotid plaque stability^18, 19^. TMAO treatment also limited macrophage M2 polarization and efferocytosis *in vitro*^19^.

The above mechanisms, involving modulation of cholesterol metabolism and inflammation, partially explain how TMAO exerts its pro-atherogenic and pro-rupture effects. We now report detailed single cell RNA-sequencing (scRNAseq) studies as the effects of TMAO on vascular cells in the context of atherosclerosis using low-density lipoprotein receptor deficient (*Ldlr−/−*) mice fed a high-cholesterol (HC) diet. Our results show that TMAO reduces the formation and increases the degradation of extracellular matrix in atherosclerotic lesions, thereby destabilizing lesions and making them more prone to rupture. In particular, we address a fibroblast-like vSMC subset, termed modulated vSMCs, that are transcriptionally distinct from contractile vSMCs. Modulated vSMCs highly express fibronectin 1 (*Fn1*), galectin 3 (*Lgals3*), and osteopontin (*Spp1*), and are present in the lesion and fibrous cap^20^. The effects of TMAO on vascular cell subtypes are poorly understood. Based on these data, we modeled both intra- and intercellular gene interactions impacted by TMAO.

## Methods

### Animals and Dietary Treatment

All animal experiments were approved by the University of California, Los Angeles (UCLA) Animal Care and Use Committee. Female low-density lipoprotein receptor deficient (*Ldlr−/−*; B6.129S7-*Ldlr^tm1Her^*/J; stock: 2207) mice were obtained from the Jackson Laboratory and maintained on a 12 hr light-dark cycle from 6 am to 6 pm. Mice were divided into 3 treatment groups: (1) Chow diet group, 14 weeks of age; (2) 0.5% cholesterol diet for 3 months, 22 weeks of age to induce lesions; (3) 0.5% cholesterol + 0.125% TMAO diet for 3 months, 22 weeks of age. The 0.125% TMAO was similar in concentration to a previous study^2^. Mouse euthanasia was performed by isoflurane overdose followed by cervical dislocation. Upon euthanasia, mice were perfused with phosphate buffered saline (PBS), and tissues were weighed and instantly frozen in liquid nitrogen. The ascending aorta, aortic arch, and thoracic aorta were isolated and prepared for scRNAseq.

### Aorta Dissection and Single Cell Dissociation

Each treatment group consisted of 3 aorta pools with n=2 mice/pool, for a total of 9 independent aorta pools from 18 mice. Aorta samples from the above groups were collected from the lesion prone areas of aorta, including the ascending aorta, aortic arch, and thoracic aorta. Aorta cell dissociation protocol was based on Wirka et al^20^. Briefly, the aorta samples were cut into <1 mm pieces and incubated in 1 ml of Hanks’ Balanced Salt solution containing 2 units of liberase and 24 units of elastase at 37°C for 1 hour. Fetal bovine serum (10% of total volume) was then added to inactivate the enzymes. The digested cells were then passed through a 70 µm cell strainer and rinsed with 4 ml PBS. After centrifugation at 300g, 4°C, for 10 min, the supernatant was discarded. The cells were then resuspended in 4 ml PBS/0.04% BSA, spun down again, and resuspended in 100 ul PBS/0.04% BSA.

### scRNAseq Library Construction and Sequencing

Approximately 16,000 aortic cells pooled from two mice within the same treatment group were used for each single cell library construction. In total, 9 independent libraries (3 libraries/diet group) were made using the 10x Genomics Chromium Next GEM Single Cell 3’ GEM, Library & Gel Bead Kit v3.1. The 9 libraries were then sequenced in one lane of NovaSeq S4 2×100bp at 2.4 billion reads.

### scRNAseq Data Preprocessing and Quality Control

Fastq files from each mouse aorta pool were individually aligned to the *mus musculus* genome assembly GRCm38 (mm10) using CellRanger version 6.0.1 (10X Genomics). CellBender was used to learn the background noise profile and differentiate cell-containing droplets from cell-free droplets^21^. A digital gene expression matrix, in which each row represents the read count of a gene and each column corresponds to a unique single cell, was generated from each library’s filtered feature-barcode datasets and was subsequently analyzed using the R package Seurat version 4.3.0^22, 23^. The number of expressed genes, number of unique molecular identifiers (UMIs), percentage of mitochondrial genes, and percentage of hemoglobin genes were considered in identifying outliers. Single cells were selected based on a threshold of between 200 and 5000 genes expressed and between 500 and 20,000 UMIs counted to reduce the presence of doublets as well as a mitochondrial gene expression cutoff of <15% and a hemoglobin gene expression cutoff of <0.1% to remove poor quality or contaminated cells. Raw transcript counts of each gene were normalized relative to the total number of read counts in that cell, and the resulting values were multiplied by 10,000 and log-transformed.

### Cell Clustering and Cell Type Annotation and Distribution

Single cells were projected onto two dimensions for visualization using Uniform Manifold Approximation and Projection (UMAP) and assigned into clusters using Louvain clustering^22, 23^. Non-parametric Wilcoxon rank sum test was used to determine cluster-specific marker genes consistent across samples and expressed in at least 10% of the cells in each cluster of interest. To resolve the identities of the cell clusters, cluster-specific marker genes were evaluated for convergence on known aorta cell type marker genes curated from previous studies, including Wirka et al., Cochain et al., Kan et al., and Kim et al^20, 24, 25, 26^. vSMCs and macrophages were further extracted to identify known subtypes within each cell type. The cell type and subtype proportions for all cell types within a diet treatment group were calculated, and significant differences in the same cell (sub)type between dietary conditions were evaluated by one-way ANOVA and Tukey’s Honest Significant Difference post-hoc test.

### Differential Gene Expression Analysis and Pathway Enrichment

Genes expressed in at least 10% of single cells for each individual cluster were considered for differential expression analysis. Non-parametric Wilcoxon rank sum test was used to compare gene expression between dietary treatment groups to identify differentially expressed genes (DEGs) in each cell cluster. We corrected for multiple testing with the Bonferroni method, and DEGs with at least a 0.1 log2 fold change in gene expression between groups and with Bonferroni-adjusted p-value <0.05 were included in downstream pathway enrichment analysis utilizing pathways from Reactome^27, 28^. However, for clusters with an insufficient number of DEGs at FDR<0.05 to identify meaningful significant pathways, DEGs with a threshold of p-value<0.01 were used to predict suggestive pathways for these cell types. Significant enrichment of pathways was based on a hypergeometric test followed by multiple testing correction with the Benjamini-Hochberg method.

### Single Cell Gene Regulatory Network (GRN) Construction

Single Cell INtegrative Gene regulatory network inference (SCING) was used to construct cell type-specific intracellular global GRNs from the scRNAseq raw counts matrix using gradient boosting^29^. SCING pseudo-bulks cells through Leiden clustering to mitigate scRNAseq gene sparsity and constructs a consensus GRN across subsamples of the data via bootstrapping. We identified highly connected subnetworks, or modules, using Leiden clustering with the final GRN. Expression profiles of GRN modules were generated using the module eigengene approach from hdWGCNA^30^. Differential module eigengenes were calculated using the Wilcoxon rank sum test with FindDMEs in hdWGCNA. Cytoscape (v. 3.8.2) was used for network visualization^31^.

### Cell-cell Communication Network Construction

CellChat was used to predict alterations in cell-cell communication in response to TMAO^32^. CellChat objects were created for HC+TMAO and HC conditions separately using endothelial cells, fibroblasts, vSMC subsets, and macrophage subsets. The total number of interactions of the inferred cell-cell communication networks from HC+TMAO and HC conditions were then compared. We also used CellChat to determine specific cell-cell signaling changes of modulated vSMCs between HC+TMAO and HC groups. To predict upstream ligands that may regulate the HC+TMAO vs. HC DEGs, we used NicheNet^33^, where modulated vSMCs were set as the receiver cell type, and the HC+TMAO vs HC modulated vSMC DEGs were inputted as the targets. We focused on vSMC DEGs involved in ECM organization, collagen formation, and apoptosis due to their significance in pathways analysis. We also carried out NicheNet analysis on macrophages, where macrophage subtypes and the HC+TMAO vs HC DEGs for each macrophage subtype were set as the targets. The sender cell types included macrophage subtypes, vSMC subtypes, endothelial cells, and fibroblasts.

### In Vitro Validation of the Direct Effect of TMAO on Key vSMC and Macrophage Genes

Human immortalized coronary artery SMCs (gift from Dr. Clint Miller, University of Virginia, Charlottesville, VA) and human primary aortic SMCs (ATCC, PCS-100-012) were cultured in SMC growth medium with premixed supplements (PromoCell, Cat# C-22062). RAW 264.7 mouse macrophage cells (ATCC, Manassas, VA) were cultured in DMEM medium supplemented with 5% FBS, 100 U/mL penicillin, and 100 ug/mL streptomycin. For TMAO treatment, cells were grown in 6-well plates overnight, followed by treatment with various concentrations of TMAO (100, 200, or 400 µM) or vehicle (PBS) in growth media for 24 hours before harvest. TMAO concentrations were selected to reflect normal physiologically relevant and elevated pathological conditions and were likewise used in our previously published work^13^. All cell culture experiments were performed in triplicate to confirm the robust direct function of TMAO. Cells were homogenized in TRIzol (Qiagen), and RNA extraction was performed according to the manufacturer’s protocol. RNA samples were reverse transcribed using a high-capacity cDNA reverse transcription kit (Applied Biosystems, Waltham, MA). qPCR was performed using the KAPA SYBR Fast qPCR kit (Kapa Biosystems, Wilmington, MA) for select target genes from the scRNAseq analysis. All qPCR primer sequences are listed in **Table S1**. Housekeeping genes included *Rpl13a* for RAW 264.7 mouse macrophage cells, *B2M* for human immortalized coronary artery SMCs, and *SRSF4* for human primary aortic SMCs. Differences in the expression level of each gene between treatment groups were determined by ANOVA.

### Immunofluorescence Staining to Validate the Effect of TMAO on Fibrous Cap Thickness In Vivo

Female *Ldlr−/−* female mice were fed HC+TMAO or HC diets for 5 months (n=5/group) to allow for the development of the fibrous cap. Aortic root was collected for fibrous cap measurements. OCT-embedded aortic root tissues were sectioned at a thickness of 10 µm with three sections per slide. Slides were fixed in 4% paraformaldehyde at room temperature, permeabilized with Triton X-100 on ice, and blocked for 1 hour at room temperature with a mixture of PBS, 3% bovine serum albumin (BSA) powder, and 5% goat serum. Slides were then incubated with transgelin (Tagln) primary antibody (Abcam, ab14106; 1:200) to stain for fibrous cap or 10x-diluted block solution as negative control. Each slide had two sections stained with the primary antibody and one section used for negative control. Goat-antirabbit 488 (Thermo Fisher Scientific, A11008; 1:500) was as the secondary antibody for immunofluorescence. Slides were washed in PBS baths between antibody steps. Slides were mounted with DAPI (Sigma Aldrich). All immunostainings were captured with Zeiss (Axioskop 2 plus, Carl Zeiss, Heidelberg, Germany) or Keyence BZ-X (Keyence, Itasca, IL) microscopes. Fibrous cap measurement was performed based on Newman et al^34^. The fibrous cap area of measurement was determined by the contiguous presence of Tagln+ cells (Tagln+ staining that was not interrupted by Tagln-DAPI+ cells). Using ImageJ, we added 15 evenly spaced points along the lumen of female *Ldlr−/−* mice fed HC+TMAO or HC diets. Lines covering the depth of the fibrous cap were drawn perpendicular to the points on the lumen, and the average of the lengths of the lines was calculated to determine the depth of the fibrous cap. Differences between groups were determined by unpaired t-test.

## Results

### scRNAseq Identification of Distinct Cell Types in the Ldlr−/− Mouse Aorta

Female *Ldlr−/−* mice (n=6/group, 22 weeks of age at euthanasia) were fed high-cholesterol (HC) or HC with TMAO (HC+TMAO) diets for 3 months, and ascending aorta, aortic arch, and thoracic aorta were collected for scRNAseq; the same aortic tissues from control female *Ldlr−/−* mice (14 weeks of age) were also collected for sequencing (**Figure 1A**). Plasma TMAO levels were significantly increased in mice fed with a HC+TMAO diet compared to HC or Chow diets (**Figure 1B**). We detected 10 main cell clusters that each displayed distinct gene expression patterns and that represented major cell types within the lesion-prone areas of the aorta (**Figure 1C-E**). Known aortic cell types recovered included vSMCs, fibroblast, macrophage, and endothelial cells. We also recovered small populations of pericytes, T cells, B cells, and neurons.

**Figure 1.**
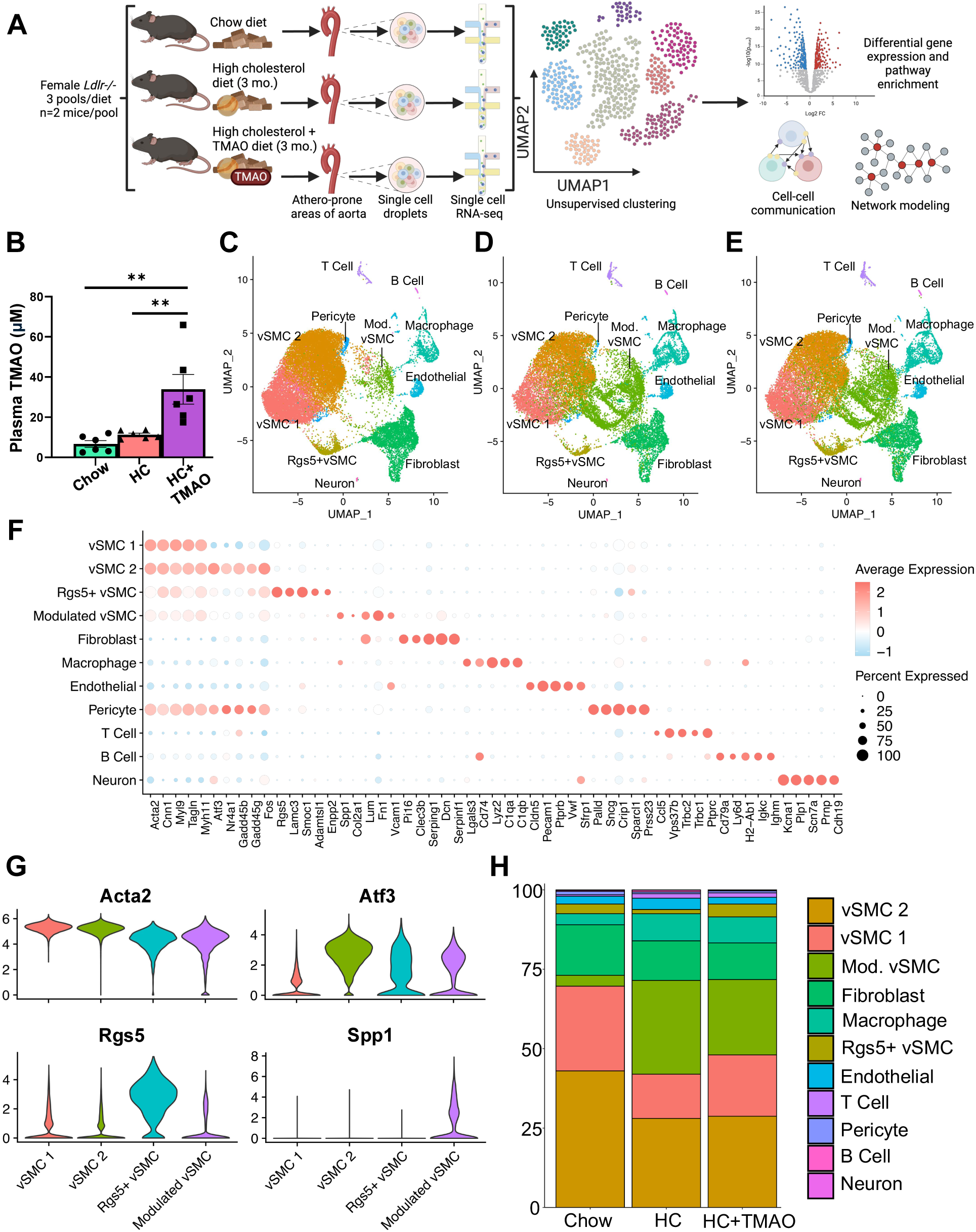
Identification of major aortic cell types and cell type-specific gene markers. **A**, Schematic diagram of the study design. Athero-prone areas of the aorta (ascending aorta, aortic arch, thoracic aorta) were collected from female *Ldlr−/−* mice on Chow, HC, or HC+TMAO diets (n=6/group), and two mouse samples from the same treatment group were pooled together, with n=3 pools/group. Chow-fed mice were 3 months of age whereas HC- and HC+TMAO-fed mice were 5 months of age with Chow diet for 2 months and switched to HC or HC+TMAO for 3 months. **B**, Plasma TMAO levels (uM) in the Chow-, HC-, and HC+TMAO-fed mice as measured by mass spectrometry (n=6/group). **C-E**, UMAP representation of cell clusters in Chow, HC, and HC+TMAO conditions, respectively. **F**, Cluster-specific expression of previously known cell type markers. **G**, Normalized expression values of top markers of vSMC subtype clusters: vSMC 1 – *Acta2*, vSMC 2 – *Atf3*, Rgs5+ vSMC – *Rgs5*, modulated vSMC – *Spp1*. **H**, Proportions of identified cell types within total cells recovered for each diet condition. Statistical significance was determined by one-way ANOVA and Tukey post-hoc test. No cell type showed significant difference in cell proportions between groups.

Subclustering of vSMCs identified four subtypes. These include two distinct classic contractile vSMC clusters, with vSMC 1 expressing higher classic contractile vSMC markers (*Acta2, Tagln, Myh11, Tpm1*) and vSMC2 expressing higher levels of *Atf3, Nr4a1, Gadd45g,* and *Gadd45b* (**Figure 1F-G**). We further identified a distinct vSMC cluster highly expressing *Rgs5* which may represent non-proliferative vSMCs that do not contribute to the fibrous cap (**Figure 1G**). Finally, we identified modulated vSMCs primarily present in the HC+TMAO and HC samples and characterized by higher expression of *Spp1, Fn1, Col2a1, Lum,* and *Lgals3* compared to other vSMC subtypes (**Figure 1F-G**). Consistent with previous findings, a small proportion of modulated vSMCs were present in the Chow condition. Other than a non-significant trend towards decreased modulated vSMCs with TMAO feeding, the single-cell distribution from HC+TMAO versus HC mice was similar (**Figure 1H**). As expected, modulated vSMCs were significantly increased with HC and HC+TMAO diets compared to Chow, which parallels the presence of lesions.

We also subclustered the macrophage cell type, revealing monocytes and resident inflammatory, foamy *Trem2*^+^, and SMC-derived macrophages (**Figure 2A-D**). SMC-derived macrophages composed 43% of the TMAO-fed macrophage population, as compared to 27% in Chow and 33% in HC groups, suggesting TMAO may promote vSMC switching towards a macrophage phenotype (**Figure 2E**).

**Figure 2.**
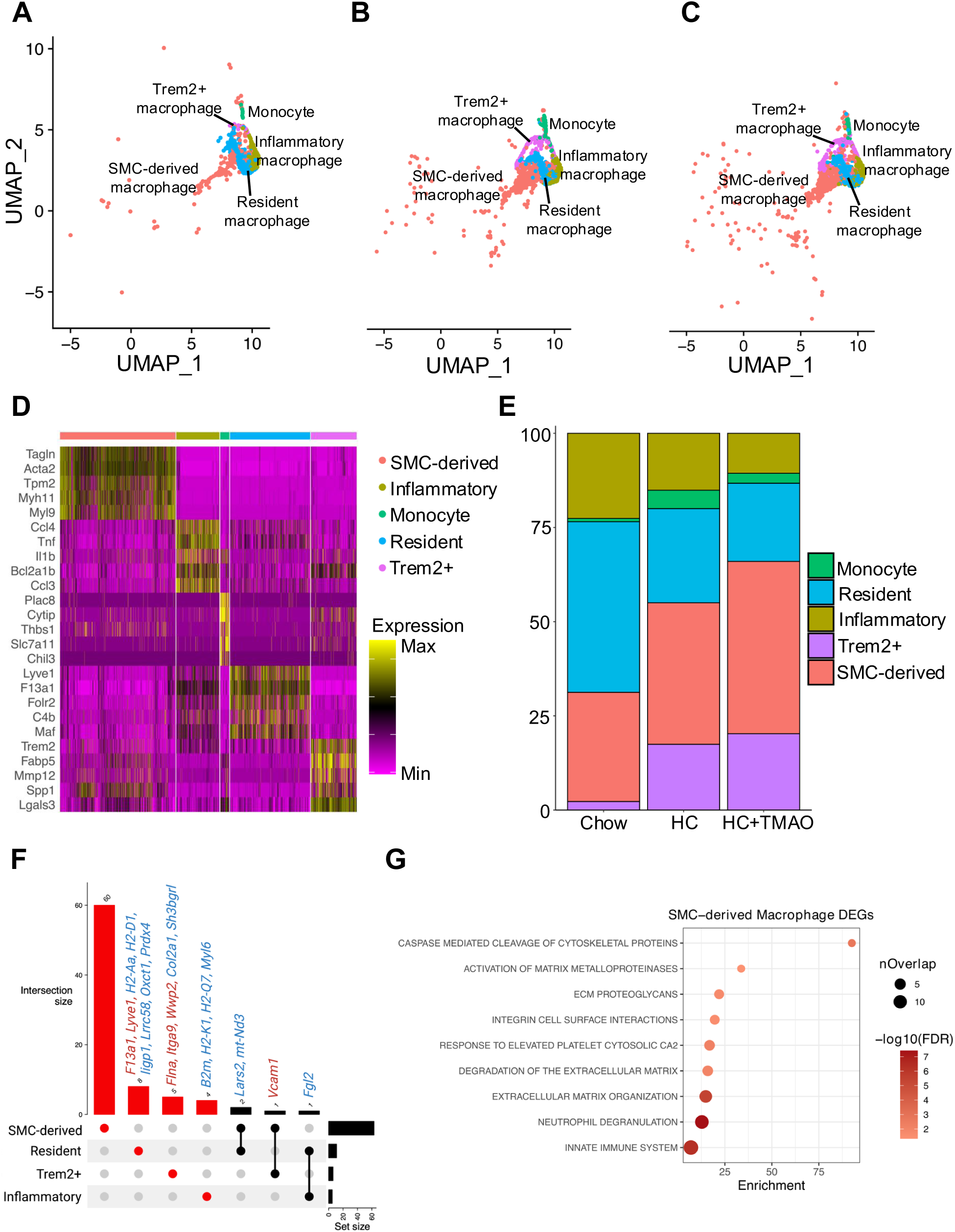
Identification of macrophage subtypes and differences in SMC-derived macrophages with TMAO. **A-C**, UMAP of macrophage subclusters in Chow, HC, and HC+TMAO conditions, respectively. **D**, Subcluster-specific expression of previously known macrophage subtype markers. **E**, Proportions of macrophage subtypes within total macrophage number recovered for each diet condition. Statistical significance was determined by one-way ANOVA and Tukey post-hoc test, and no cell type showed significant difference in proportions between groups. **F**, Shared and cell-type specific DEGs (FDR<0.05) induced by TMAO feeding in macrophage subtypes. DEGs unique to a cell type are highlighted in red in the upset plot and histogram, and shared DEGs are indicated in black. The histogram above each plot indicates the DEG counts for each category. DEG direction is indicated by the color of the gene name: red – upregulated, blue – downregulated. **G**, Top representative pathways from Reactome enriched for SMC-derived macrophage DEGs (FDR<0.05).

Overall, our scRNAseq analysis revealed all expected aortic cell types and subtypes, as well as the increases in disease-associated vSMCs and macrophages in atherosclerosis conditions. Importantly, compared to HC only, TMAO+HC exhibited trends towards decreased proportions of atheroprotective vSMCs and increased proportions of SMC-derived macrophages.

### Confirmation of Key Atherogenic Genes and Pathways Between HC and Chow Groups

We first identified DEGs between HC and Chow groups, confirming the atherogenic effects of HC in our scRNAseq data (**Figure S1A, Table S2**). For example, upregulated DEGs in modulated vSMCs (FDR<0.05) were significantly enriched for ECM organization, programmed cell death, and platelet activation and aggregation (**Figure S1B, Table S5**). Modulated vSMC downregulated DEGs were significantly enriched for muscle contraction, supporting that vSMCs phenotypically switch from classic contractile towards a synthetic phenotype with atherosclerosis progression (**Figure S1C, Table S6**).

### TMAO Impacts Shared and Cell Type-specific Genes and Biological Pathways

To understand the ubiquitous and cell type-specific impact of elevated TMAO in lesions, we identified shared and cell type-specific DEGs between HC+TMAO and HC (**Table S3**). Across cell types, various mitochondria-encoded genes (e.g., *mt-Nd3*, *mt-Atp8*, and *mt-Nd2*) were downregulated with the addition of TMAO (**Figure 3A**), suggesting potential mitochondrial dysfunction and increased oxidative stress across cell types with TMAO. Mitochondrial dysfunction can trigger various cellular stress responses, including apoptosis. In support of a direct effect of TMAO, human coronary artery SMCs treated with varying concentrations of TMAO exhibited downregulation of *ATP8* and *ND3* (**Figure 3D**). Because TMAO supplementation in the diet can affect TMAO-TMA conversion via *Fmo2* (TMAO to TMA) and *Fmo3* (TMA to TMAO), we found *Fmo2* to be upregulated in six cell types and *Fmo3* to be upregulated in three vSMC subtypes.

**Figure 3.**
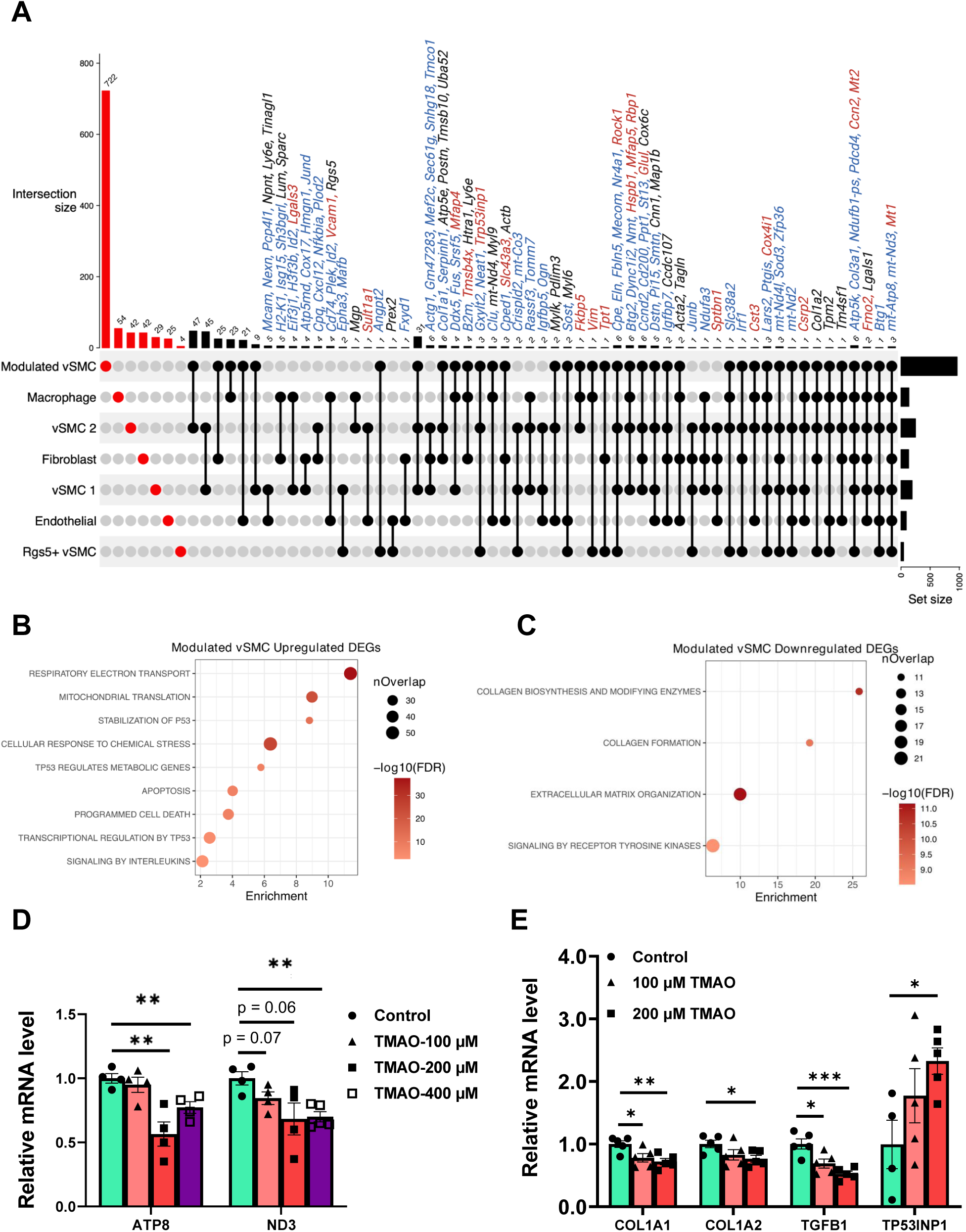
Shared and cell-type specific DEGs induced by TMAO in major aortic cell types. **A,** Shared and cell-type specific DEGs (FDR<0.05) induced by TMAO feeding in major aortic cell types recovered in scRNAseq. DEGs unique to a cell type are highlighted in red in the upset plot and histogram, and shared DEGs are indicated in black. The histogram above each plot indicates the DEG counts for each category. DEG direction is indicated by the color of the gene name: red – upregulated in HC+TMAO vs HC, blue – downregulated in HC+TMAO vs HC, black – cell type-dependent. **B-C,** Top representative pathways from Reactome enriched for upregulated and downregulated modulated vSMC DEGs (FDR<0.05), respectively. **D,** Mitochondrial genes *ATP8* and *ND3* are downregulated in immortalized human aortic smooth muscle cells when treated with TMAO for 24 hours. **E,** ECM-related genes *COL1A1, COL1A2*, and *TGFB1* were downregulated whereas p53 proapoptotic target gene *TP53INP1* was upregulated in primary human coronary artery smooth muscle cells when treated with TMAO for 24 hours. One-way ANOVA with Tukey’s post hoc test was used to determine significance.

Among all cell types, modulated vSMCs exhibited the greatest number of DEGs (n=962) that are affected by TMAO, and the majority of DEGs were specific (n=722) to this atherosclerosis-associated vSMC subtype (**Figure 3A**). Upregulated modulated vSMC DEGs were enriched for respiratory electron transport (nuclear-encoded cyclooxygenase and NADH: ubiquinone gene families), apoptosis, and P53-related pathways (**Figure 3B, Table S7**). *Trp53inp1*, a proapoptotic P53 target gene, was upregulated in modulated vSMCs *in vivo*, and we confirmed a direct effect of TMAO on this gene in primary human SMCs (**Figure 3E**). Previous work has shown that adenoviral-mediated expression of p53 in mouse atherosclerotic lesions resulted in vSMC apoptosis, cap thinning, and increased vulnerability to rupture^35^. Downregulated modulated vSMC DEGs were enriched for collagen biosynthesis and formation, extracellular matrix (ECM) organization, and signaling by receptor tyrosine kinases (**Figure 3D, Table S8**). We further confirmed the direct effect of TMAO on downregulating key collagen genes *COL1A1* and *COL1A2* as well as regulator *TGFB1* in primary human aortic SMCs (**Figure 3E**). As phenotypically switched vSMCs contribute to lesion composition and largely compose the fibrous cap, reduced collagen and dysregulated ECM organization by TMAO may result in a more vulnerable fibrous cap. These results suggest that TMAO decreases collagen and ECM content and promotes apoptosis of phenotypically switched vSMCs, events that exacerbate lesion cap instability.

Macrophage subtypes also exhibited cell-type specific transcriptomic changes induced by TMAO feeding (**Figure 2F, Table S4**). Specifically, SMC-derived macrophage DEGs were significantly enriched for ECM- and immune-related pathways, activation of matrix metalloproteinases (MMPs), and caspase-mediated cleavage of cytoskeletal proteins, a hallmark of apoptosis (**Figure 2G, Table S9**). Thinning of the fibrous cap is also attributed to release of MMPs that degrade the ECM by vSMCs or SMC-derived macrophages^36^. Additionally, *Trem2^+^* foamy macrophage DEGs (p-value<0.01) were enriched for ECM proteoglycans, integrin cell surface interactions, ECM organization, and autophagy (**Figure S2A, Table S9**). DEGs from the remaining macrophage subtypes were enriched for immune and inflammatory pathways, such as mitophagy, antigen processing cross presentation, and interferon and cytokine signaling in inflammatory macrophages (**Figure S2B, Table S9**), C type lectin receptors, death receptor signaling, interleukin and cytokine signaling in monocytes (**Figure S2C, Table S9**), and interferon signaling, TGFβ signaling, and antigen processing and presentation in resident macrophages (**Figure S2D, Table S9**).

### Within-cell-type Regulators of TMAO DEGs and Pathways in vSMCs

To complement the DEG and pathway analysis and elucidate intracellular regulatory cascades induced by TMAO feeding, we constructed cell type-specific directed gene regulatory networks (GRNs) using an unbiased gradient boosting method SCING. We identified subnetworks (termed modules) associated with TMAO in each cell type. For modulated vSMCs, modules significantly associated with TMAO confirmed the ECM and apoptosis pathways revealed above through M11 (ECM organization and biological adhesion) and M19 (regulation of cell death), and additionally we identified M4 (translation initiation and protein transport) and M1 and M2 (electron transport chain and mitochondrion organization) (**Figure 4A**). We further retrieved the top interconnected genes within each module as potential regulators as they signify centrality and importance within the network structure. For example, *Tnn*, *Col11a2*, and *Col9a3* were potential regulators of SCING module M11, enriched for ECM organization and biological adhesion (**Figure 4B**). *Gpnmb*, *Mdk*, and *Ccl19* were regulators of the cell death module M19 (**Figure 4C**). Also, this cell death regulation GRN included three known human CAD genome-wide association study (GWAS) hits: *Fndc1*, *Mras*, and *Lipa*. For the inflammation module M13, we identified *Tyrobp*, *C1qa*, and *Cxcl2* as potential regulators (**Figure 4D**).

**Figure 4.**
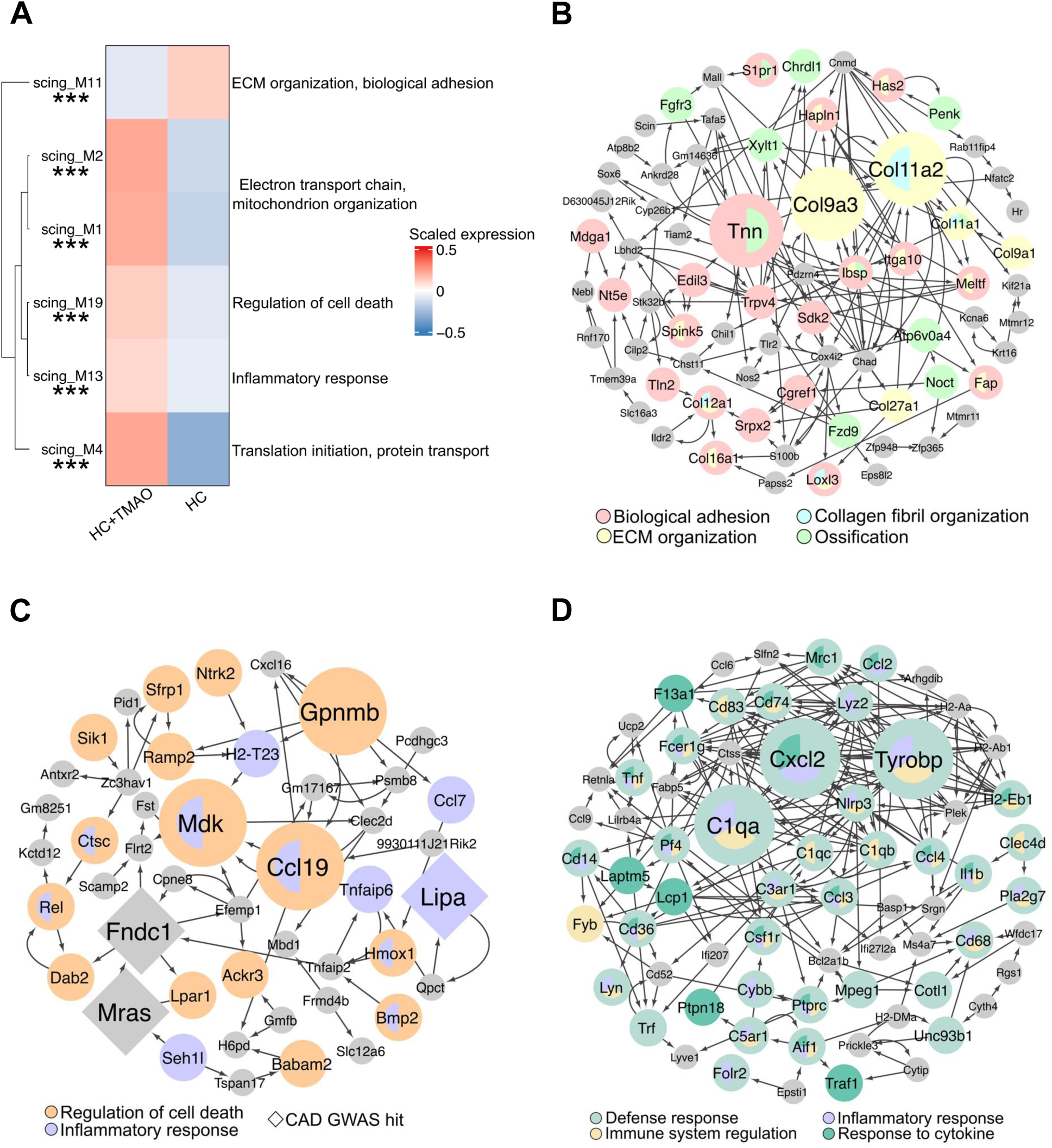
Modulated vSMC gene regulatory networks significantly associated with TMAO in the diet. **A**, Top SCING gene regulatory network modules of the modulated vSMCs significantly associated with TMAO feeding. The top biological pathways enriched for the genes in each network are listed. **B-D**, Network structures of directed SCING modules M11, M19, and M13, respectively. The direction of the interaction is denoted by arrows in the networks. The top 3 most interconnected nodes involved in the relevant pathways for each network and known human coronary artery disease GWAS hits are highlighted by larger size. GWAS hits are also denoted by the diamond shape.

### TMAO Impacts Cell-cell Communication and External Signaling Regulators Targeting Modulated vSMCs and Macrophage Subtypes

To further understand the atherosclerotic microenvironment impacted by TMAO, we predicted cell-cell communications involving the major cell types within the lesion. Ligand-receptor based CellChat analysis indicated that TMAO increased modulated vSMC-endothelial and endothelial-endothelial interactions and enhanced outgoing signals from *Trem2^+^* macrophage and SMC-derived macrophage (**Figure 5A-B**). For modulated vSMCs, collagen, NCAM, and laminin signaling which are all related to ECM organization, were predicted to decrease as both incoming and outgoing interactions, whereas SPP1 signaling was predicted to increase as an incoming and outgoing signal (**Figure 5C**). *Spp1* was significantly upregulated in modulated vSMCs with TMAO feeding in our scRNAseq (**Figure S3A**) and is a well-known biomarker elevated in atherosclerotic patients that can track plaque severity and cardiovascular event mortality^37, 38^.

**Figure 5.**
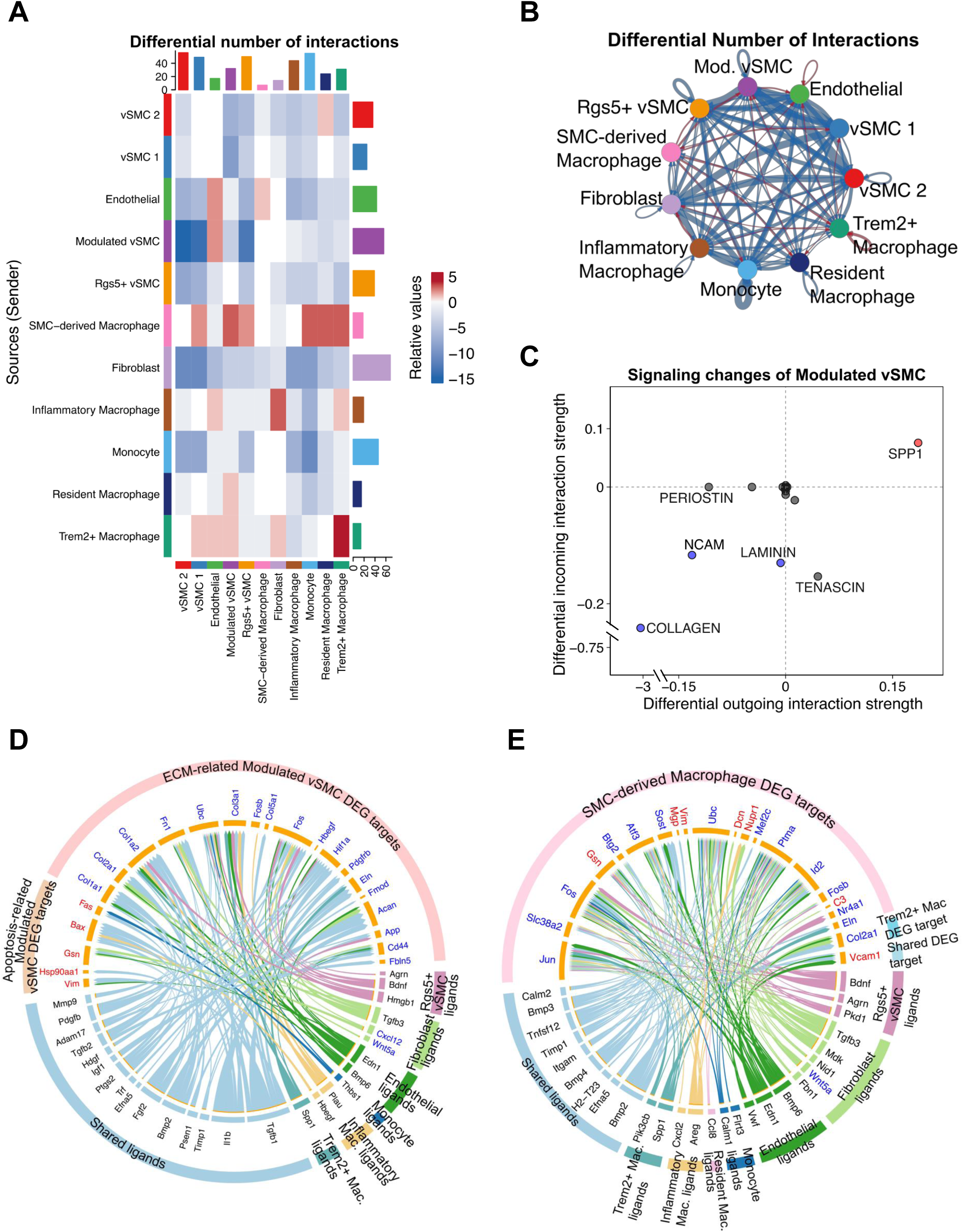
TMAO alters cell-cell communications between major cell types involved in atherosclerosis progression. **A**, Heatmap of predicted differential number of interactions between sender cells (rows) and receiver cells (columns) comparing HC+TMAO and HC by CellChat. The top-colored bar plot shows the total incoming signaling change for a cell type while the right colored bar plot shows the total outgoing signaling change in the number of signaling interactions between HC+TMAO and HC for each cell type. In the heatmap, red represents increased signaling with TMAO feeding, and blue represents decreased signaling. **B**, Network visualization of predicted differential number of interactions in the cell-cell communication network between HC+TMAO and HC by CellChat. The color of the network edges represents increased (red) or decreased (blue) signaling with TMAO feeding. **C**, Specific incoming and outgoing signaling pathways predicted to change in modulated vSMCs between HC+TMAO and HC by CellChat. Signaling pathways increased and decreased as both incoming and outgoing signals are highlighted in red and blue, respectively. **D**, Top ligands predicted by NicheNet to target the modulated vSMC DEGs involved in apoptosis or ECM organization. **E**, Top ligands predicted by NicheNet to target the SMC-derived macrophage and *Trem2^+^* macrophage DEGs. In D-E, DEG direction is indicated by red (upregulated with TMAO) or blue (downregulated with TMAO).

We next identified external signaling regulators of modulated vSMC DEGs involved in the upregulation of apoptosis and downregulation of ECM organization using NicheNet. Apoptosis-related DEGs in modulated vSMCs were predicted to be regulated by various ligands from multiple cell types, such as *Tgfb1*, *Ptgs2*, *Agrn*, *Efna*, *Adam17*, and *Trf,* and by inflammatory macrophage ligand *Hbegf*, monocyte *Thbs1*, and endothelial ligand *Edn1* (**Figure 5D**). ECM-related DEGs in modulated vSMCs were predicted to be regulated by *Trem2^+^*macrophage ligand *Spp1*, inflammatory macrophage ligand *Plau*, endothelial ligands *Edn1* and *Bmp6*, and various ligands shared across cell types, including *Tgfb1*, *Il1b*, *Timp1*, and *Mmp9* (**Figure 5D**). *Tgfb1*, one of the master regulators of ECM molecule secretion, was predicted to be a shared ligand between monocytes and inflammatory macrophages to regulate ECM genes in modulated vSMCs. We confirmed the direct effect of TMAO on significantly downregulating *Tgfb1* in RAW cells (**Figure S3B**). We further predicted ligands that target DEGs of macrophage subtypes, revealing potential regulatory ligands from endothelial cells (e.g., *Edn1*, *Bmp6*) and from other macrophage subtypes (e.g. *Bmp2*, *Spp1*) (**Figure 5E**).

### TMAO reduces the fibrous cap size in atherosclerotic lesions

As we have identified apoptosis and ECM organization as key up- and downregulated processes, respectively, in modulated vSMCs with TMAO feeding, and ECM remodeling signals from various macrophage subtypes to modulated vSMCs, we hypothesized that TMAO feeding will reduce the fibrous cap to promote plaque instability under TMAO treatment. We therefore examined the fibrous caps of *Ldlr−/−* female mice fed HC+TMAO or HC diets for 5 months. Thinner fibrous caps overlaying the atherosclerotic lesion implicate plaque vulnerability and increased likelihood of rupture, so we employed Tagln immunostaining to identify and measure thickness of the cap. We utilized depth of contiguous Tagln^+^ cells from the lumen, representing the fibrous cap, as a marker of plaque stability in aortic root lesions. Mice fed the HC+TMAO diet exhibited significantly decreased fibrous cap thickness compared to mice fed the HC diet (**Figure 6A-D**).

**Figure 6.**
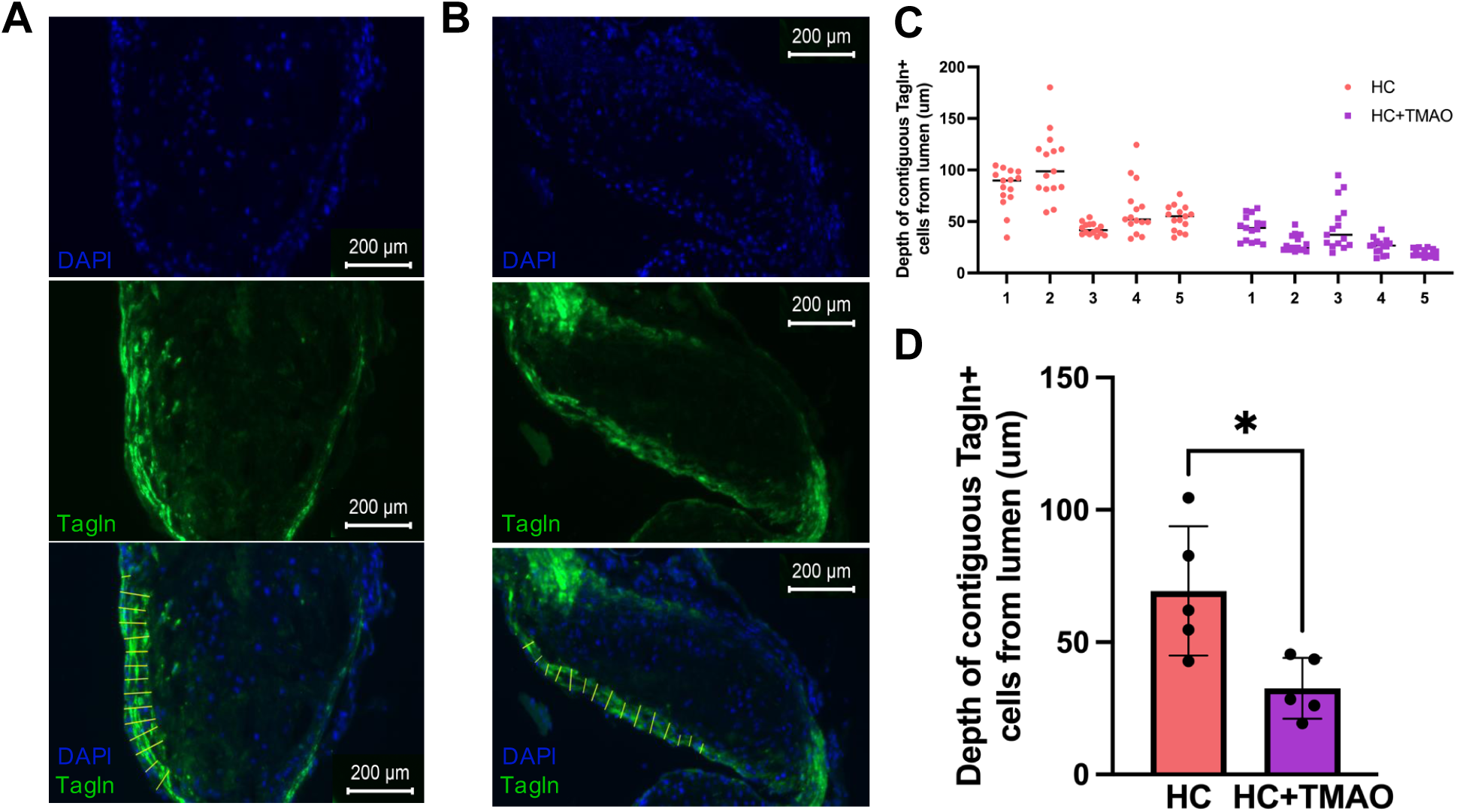
HC+TMAO vs HC lesion fibrous cap staining and quantification. **A**, Representative immunofluorescence staining of aortic root lesions and the fibrous cap in the HC group. **B**, Representative immunofluorescence staining of aortic root lesions and the fibrous cap in the HC+TMAO group. In **A-B**: DAPI (top, blue), Tagln (middle, green), and merge (bottom). **C**, Depth of contiguous Tagln+ cells from lumen (µm) measurements for each mouse (15 evenly spaced measurements/mouse, n=5 mice/group). **D**, Statistical analysis of HC+TMAO vs HC fibrous cap measurements (n=5 mice/group) by t-test. *p<0.05.

## Discussion

To our knowledge, this is the first study regarding TMAO to account for the cellular heterogeneity of the lesion microenvironment and to characterize the effect of TMAO on the gene expression signatures of *in vivo* disease-specific vascular cell types. Our studies highlight the particular importance of modulated vSMCs and SMC-derived macrophages, two key atherosclerosis-associated cell types, in TMAO actions. Our findings from differential gene expression analysis, intracellular gene regulatory network modeling, and cell-cell communication predictions converged to reveal that TMAO upregulates apoptotic gene signatures and downregulates ECM organization and collagen formation gene signatures in atheroprotective modulated vSMCs. The direct effect of TMAO on these cell types and pathways were confirmed via *in vitro* exposure studies in vSMC and macrophage cell lines. As key pathways known to contribute to plaque instability include decreased collagen production at the fibrous cap by phenotypically switched vSMCs and increased collagen degradation via macrophage-released MMPs, our scRNAseq results suggest TMAO may promote plaque instability and likelihood of rupture, which is supported by the thinner fibrous caps under TMAO treatment (**Figure 6E**).

In addition to standard DEG and pathway analysis which highlighted cell type specific alterations such as downregulation of ECM and increase of apoptosis in modulated vSMCs, our within- and between-cell-type network analyses offered comprehensive insights into the regulatory cascades within and across cell types via various regulators. This included decreased collagen and laminin signaling in modulated vSMCs, increased *Spp1* signaling in modulated vSMCs, and predicted *Spp1* signaling from *Trem2^+^* macrophages targeting modulated vSMC ECM DEGs. Interestingly, the cell type-specific GRNs also highlighted human coronary artery disease GWAS hits *LIPA*, *MRAS*, and *FNDC1*^39, 40, 41^ present in a modulated vSMC subnetwork enriched for regulation of cell death and inflammatory response^39^. This mechanistic interpretation of the GWAS variants and further highlights their cellular context and connection to TMAO risk.

It is important to note that we identified various ECM and collagen genes as significantly downregulated in modulated vSMCs by TMAO. However, a previous study found no significant impact of TMAO on plaque burden or histological features, including collagen, when feeding *Ldlr−/−* or *ApoE−/−* mice a high-versus low-choline high-fat, high cholesterol diet^18^. This difference in findings may be attributed to differences in diets, mouse age, and the experimental measure used to characterize plaque instability (i.e. scRNAseq gene expression vs histology). However, our findings align with other previous work^3, 42^. To our knowledge, our study is the first to show that TMAO to promotes apoptosis in the context of atherosclerosis and to identify a subset of athero-protective modulated vSMCs to be most susceptible to TMAO. This agrees with a recent study revealing a TMAO to abdominal aortic aneurysm link via upregulation of endoplasmic reticulum stress and apoptosis genes in aortic vSMCs^3^. Also, although platelets could not be captured in our aorta scRNAseq, we observed a potential downstream effect of platelet response to TMAO as the endothelial cell DEGs in our data were significantly enriched for platelet activation signaling and aggregation. Previously, TMAO has also been shown to directly act on platelets to promote platelet hyperresponsiveness and a prothrombotic phenotype by altering calcium release from intracellular stores^42^.

While our findings elucidated the impact of TMAO across vascular cell types, we also acknowledge the limitations of our study. First, our study focuses on gene expression signatures via scRNAseq and *in vitro* gene expression quantification. Additional validation at protein and functional levels is warranted. In addition, certain cell types recovered were low in population number, such as T cells and B cells, thereby limiting the power to perform differential gene expression analysis. Further, here we focused on measuring the thickness of the fibrous cap via Tagln immunostaining of vSMCs that constitute the cap, but additional plaque instability measures such as markers of inflammation, platelet activation, and intraplaque hemorrhage could be examined.

In summary, our findings provide molecular and phenotypic evidence that TMAO may contribute to plaque instability through regulating EMC formation and degradation in cell subtypes that arise with atherosclerosis progression, such as modulated vSMCs and SMC-derived macrophages.

## Acknowledgements

We thank the UCLA Technology Center for Genomics and Bioinformatics for the sequencing of our single cell data. We thank Dr. Brian Kim for sharing his expertise in single cell protocols. We thank Dr. Clint Miller for kindly providing the immortalized human coronary artery vSMCs. We thank the laboratories of Drs. Thomas Vallim, Elizabeth Tarling, and Peter Tontonoz for use of their microscopy facilities.

## Data Availability

All data and materials have been made publicly available at the Gene Expression Omnibus repository and can be accessed with the series record GSE290376.

## Nonstandard Abbreviations and Acronyms

CAD: coronary artery disease
DEG: differentially expressed gene
ECM: extracellular matrix
GRN: gene regulatory network
Ldlr: low density lipoprotein receptor
scRNAseq: single cell RNA sequencing
TMA: trimethylamine
TMAO: trimethylamine-N-oxide
UMAP: uniform manifold approximation and projection
vSMC: vascular smooth muscle cell

## Supplemental Material

Figures S1-S3

Table S1-S9

Major Resources Table

---

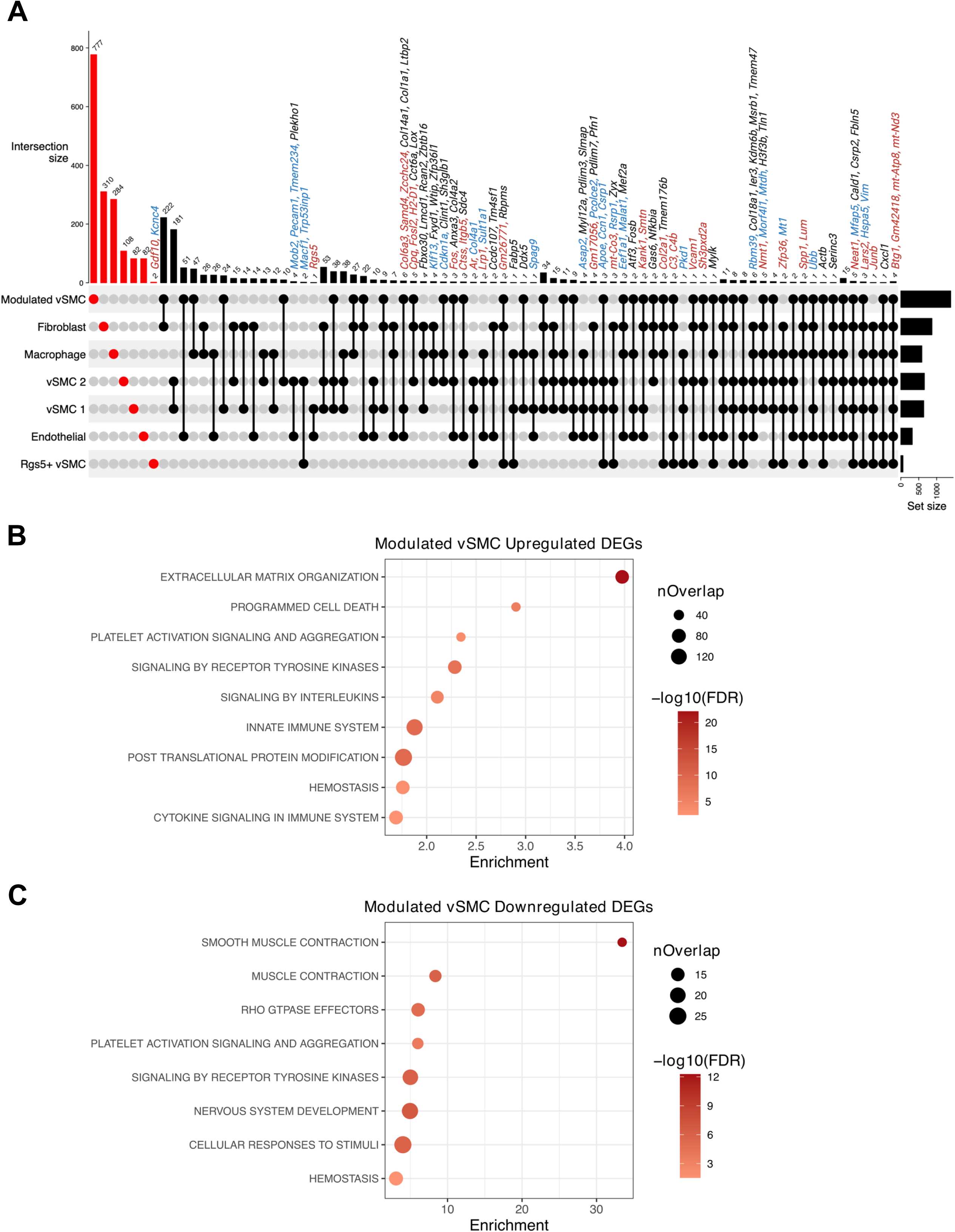

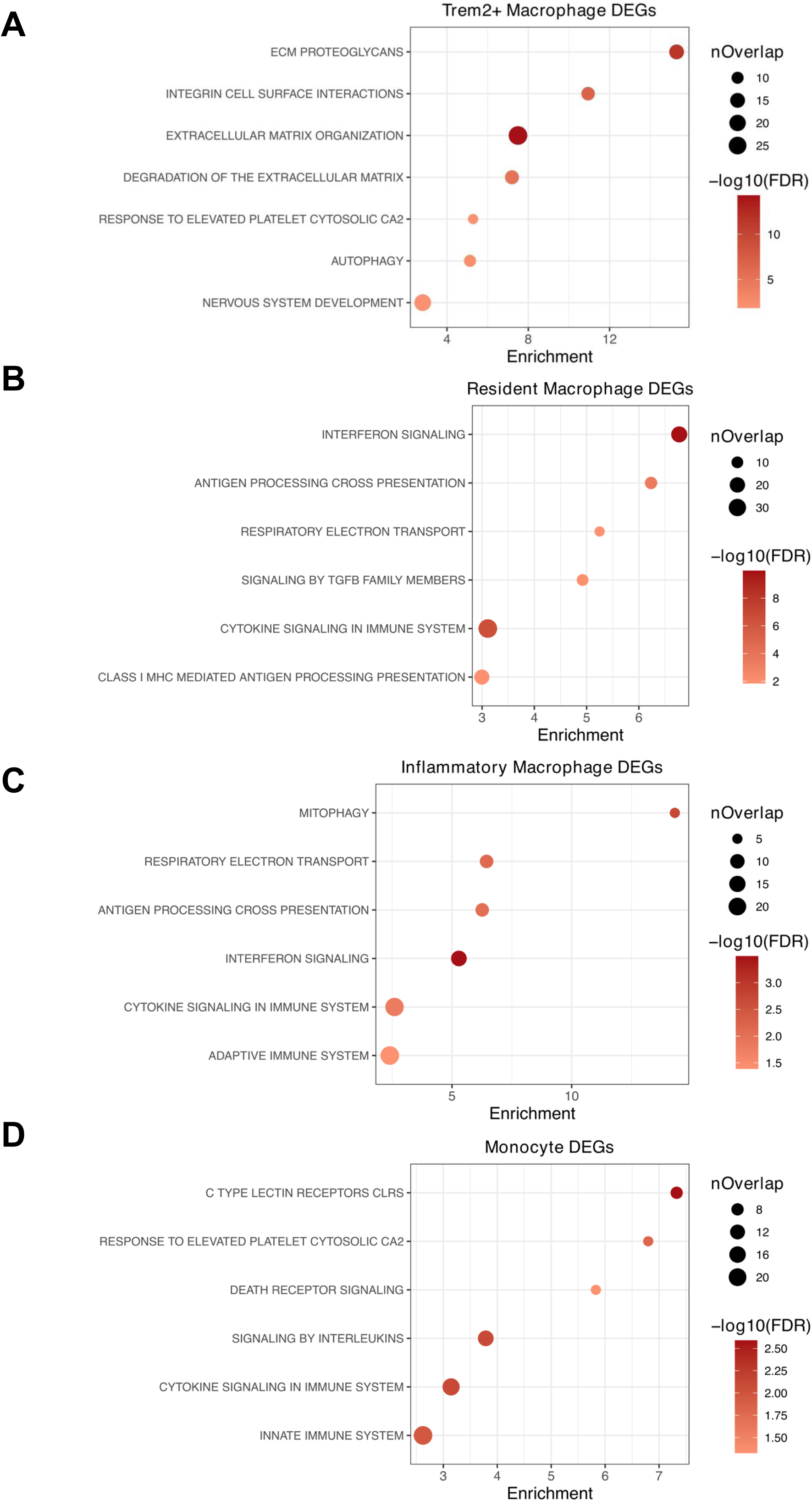

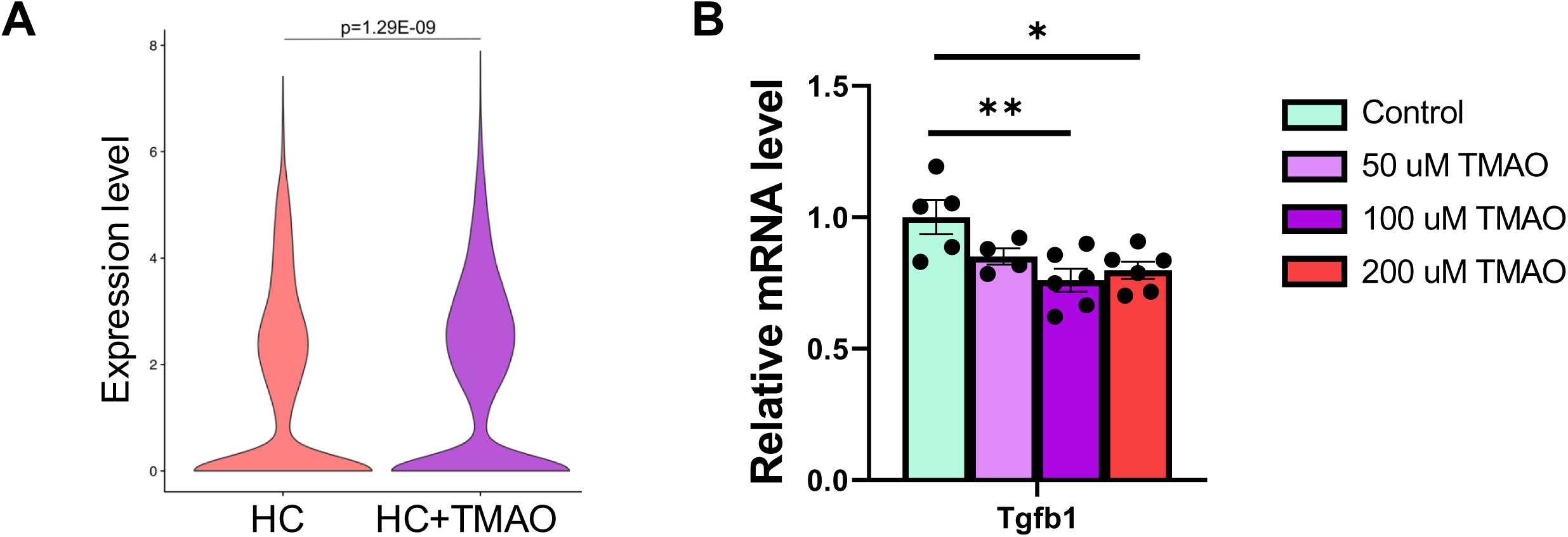

