## Supplementary material for "Trimethylamine-N-oxide affects cell type-specific pathways and networks in mouse aorta to promote atherosclerotic plaque vulnerability": Fig. S1-S3, Table S1

**Figure S1. Shared and cell-type specific DEGs induced by high-cholesterol (HC) in major aortic cell types. A**, Shared and cell-type specific DEGs (FDR<0.05) induced by HC feeding in major aortic cell types recovered in scRNAseq. DEGs unique to a cell type are highlighted in red in the upset plot and histogram, and shared DEGs are indicated in black. The histogram above each plot indicates the DEG counts for each category. DEG direction is indicated by the color of the gene name: red – upregulated with HC feeding vs Chow, blue – downregulated with HC feeding vs Chow, black – cell type-dependent. Statistical significance was determined by Wilcoxon rank sum test. P-values were adjusted by the Bonferroni method. **B-C**, Top representative pathways from Reactome enriched for upregulated and downregulated modulated vSMC DEGs (FDR<0.05), respectively. Statistical significance was determined by a hypergeometric test followed by multiple testing correction with the Benjamini-Hochberg method.

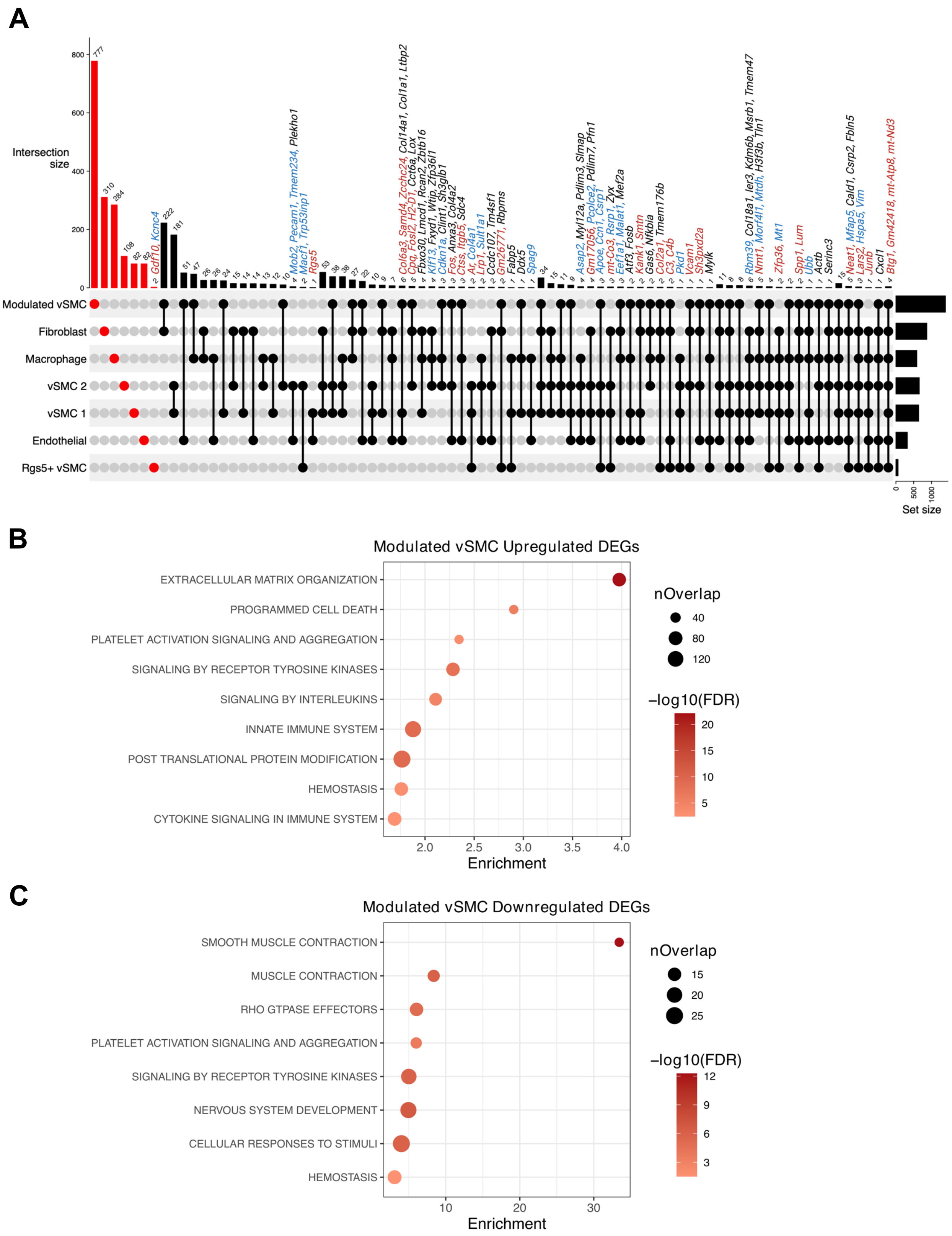
A
Intersection size
Modulated vSMC
Fibroblast
Macrophage
vSMC 2
vSMC 1
Endothelial
Rgs5+ vSMC
Set size
Gdf10, Kcnc4
Mob2, Pecam1, Tmem234, Plekho1
Macf1, Trp53inp1
Rgs5
Col6a3, Samd4, Zcchc24, Col14a1, Col1a1, Ltbp2
Cpq, Fosl2, H2-D1, Cct6a, Lox
Fbxo30, Lmcd1, Rcan2, Zbtb16
Klf13, Fxyd1, Wtip, Zfp36l1
Cdkn1a, Clint1, Sh3glb1
Fos, Anxa3, Col4a2
Ctss, Itgb5, Sdc4
Ar, Col4a1
Lrp1, Sult1a1
Cdc107, Tm4sf1
Gm26771, Rbpms
Fabp5
Ddx5
Spag9
Asap2, Myl12a, Pdlim3, Slmap
Gm17056, Pcolce2, Pdlim7, Pfn1
Apoe, Ccn1, Csrp1
mt-Co3, Rsrp1, Zyx
Eef1a1, Malat1, Mef2a
Atf3, Fosb
Kank1, Smtn
Gas6, Nfkbia
Col2a1, Tmem176b
C3, C4b
Pkd1
Vcam1
Sh3pxd2a
Mylk
Rbm39, Col18a1, Ier3, Kdm6b, Msrb1, Tmem47
Nmt1, Morf4l1, Mtdh, H3f3b, Tln1
Zfp36, Mt1
Spp1, Lum
Ubb
Actb
Serinc3
Neat1, Mfap5, Cald1, Csrp2, Fbln5
Lars2, Hspa5, Vim
Junb
Cxcl1
Btg1, Gm42418, mt-Atp8, mt-Nd3
B
Modulated vSMC Upregulated DEGs
EXTRACELLULAR MATRIX ORGANIZATION
PROGRAMMED CELL DEATH
PLATELET ACTIVATION SIGNALING AND AGGREGATION
SIGNALING BY RECEPTOR TYROSINE KINASES
SIGNALING BY INTERLEUKINS
INNATE IMMUNE SYSTEM
POST TRANSLATIONAL PROTEIN MODIFICATION
HEMOSTASIS
CYTOKINE SIGNALING IN IMMUNE SYSTEM
Enrichment
nOverlap
−log10(FDR)
C
Modulated vSMC Downregulated DEGs
SMOOTH MUSCLE CONTRACTION
MUSCLE CONTRACTION
RHO GTPASE EFFECTORS
PLATELET ACTIVATION SIGNALING AND AGGREGATION
SIGNALING BY RECEPTOR TYROSINE KINASES
NERVOUS SYSTEM DEVELOPMENT
CELLULAR RESPONSES TO STIMULI
HEMOSTASIS
Enrichment
nOverlap
−log10(FDR)

**Figure S2. Biological pathways impacted by TMAO feeding in macrophage subtypes. A-D**, Top suggestive pathways from Reactome enriched for Trem2+ macrophage, inflammatory macrophage, monocyte, and resident macrophage DEGs (HC+TMAO vs HC; p-value<0.01), respectively.

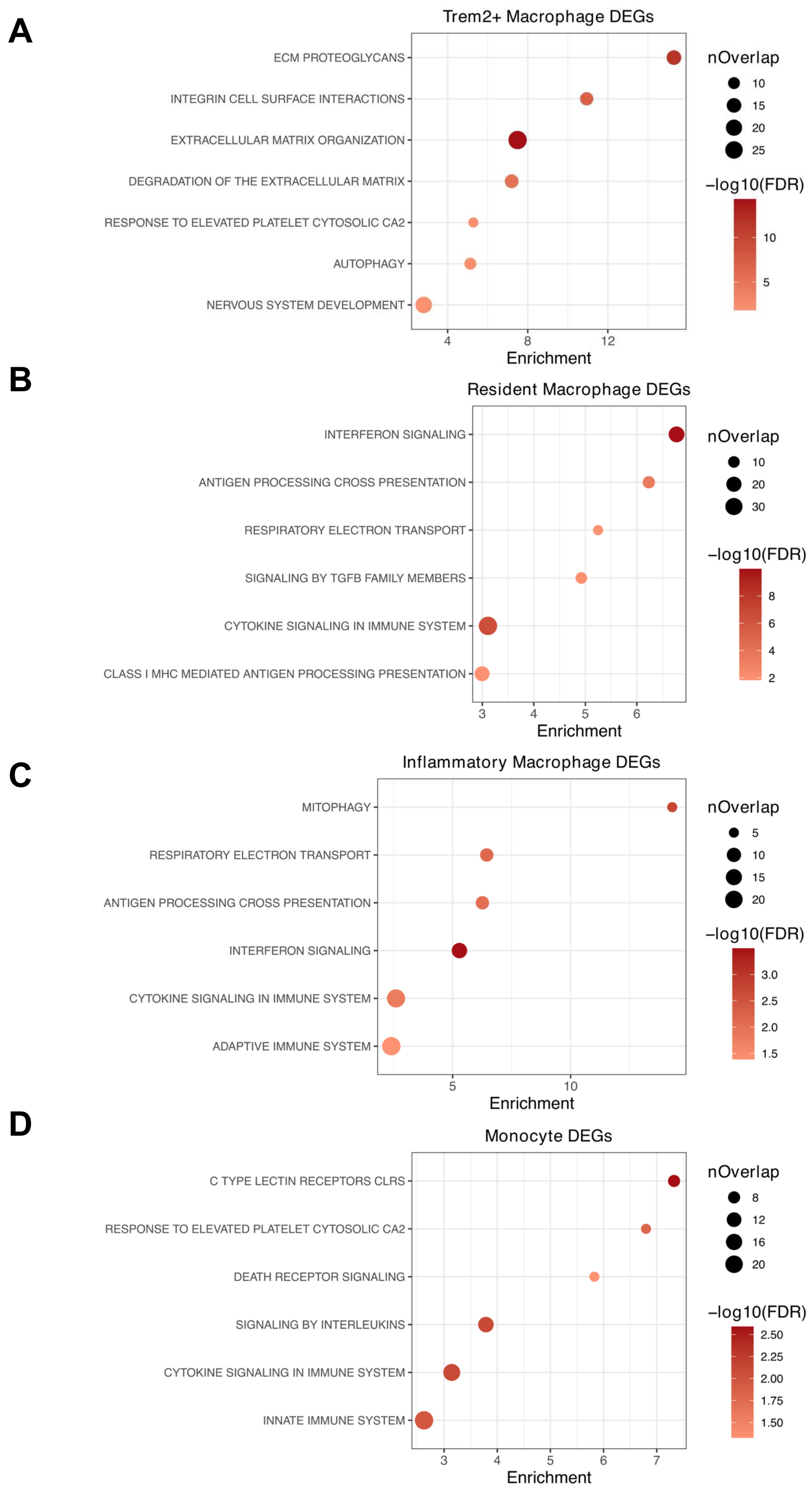

A
Trem2+ Macrophage DEGs
ECM PROTEOGLYCANS
INTEGRIN CELL SURFACE INTERACTIONS
EXTRACELLULAR MATRIX ORGANIZATION
DEGRADATION OF THE EXTRACELLULAR MATRIX
RESPONSE TO ELEVATED PLATELET CYTOSOLIC CA2
AUTOPHAGY
NERVOUS SYSTEM DEVELOPMENT
4
8
12
Enrichment
nOverlap
10
15
20
25
−log10(FDR)
10
5
B
Resident Macrophage DEGs
INTERFERON SIGNALING
ANTIGEN PROCESSING CROSS PRESENTATION
RESPIRATORY ELECTRON TRANSPORT
SIGNALING BY TGFB FAMILY MEMBERS
CYTOKINE SIGNALING IN IMMUNE SYSTEM
CLASS I MHC MEDIATED ANTIGEN PROCESSING PRESENTATION
3
4
5
6
Enrichment
nOverlap
10
20
30
−log10(FDR)
8
6
4
2
C
Inflammatory Macrophage DEGs
MITOPHAGY
RESPIRATORY ELECTRON TRANSPORT
ANTIGEN PROCESSING CROSS PRESENTATION
INTERFERON SIGNALING
CYTOKINE SIGNALING IN IMMUNE SYSTEM
ADAPTIVE IMMUNE SYSTEM
5
10
Enrichment
nOverlap
5
10
15
20
−log10(FDR)
3.0
2.5
2.0
1.5
D
Monocyte DEGs
C TYPE LECTIN RECEPTORS CLRS
RESPONSE TO ELEVATED PLATELET CYTOSOLIC CA2
DEATH RECEPTOR SIGNALING
SIGNALING BY INTERLEUKINS
CYTOKINE SIGNALING IN IMMUNE SYSTEM
INNATE IMMUNE SYSTEM
3
4
5
6
7
Enrichment
nOverlap
8
12
16
20
−log10(FDR)
2.50
2.25
2.00
1.75
1.50

**Figure S3. *Spp1* and *Tgfb1* gene expression in scRNAseq and RAW cells, respectively. A**, *Spp1* gene expression in scRNAseq data. Statistical significance was determined by non-parametric Wilcoxon rank sum test, and the p-value was adjusted by the Bonferroni method. **B**, *Tgfb1* gene expression in mouse RAW cells with TMAO treatment. RAW 264.7 cells were treated with vehicle (control), 50 μM TMAO, 100 μM TMAO, or 200 μM TMAO for 24 hours, and *Tgfb1* gene expression levels were measured via qPCR. Statistical significance was determined by t-test.

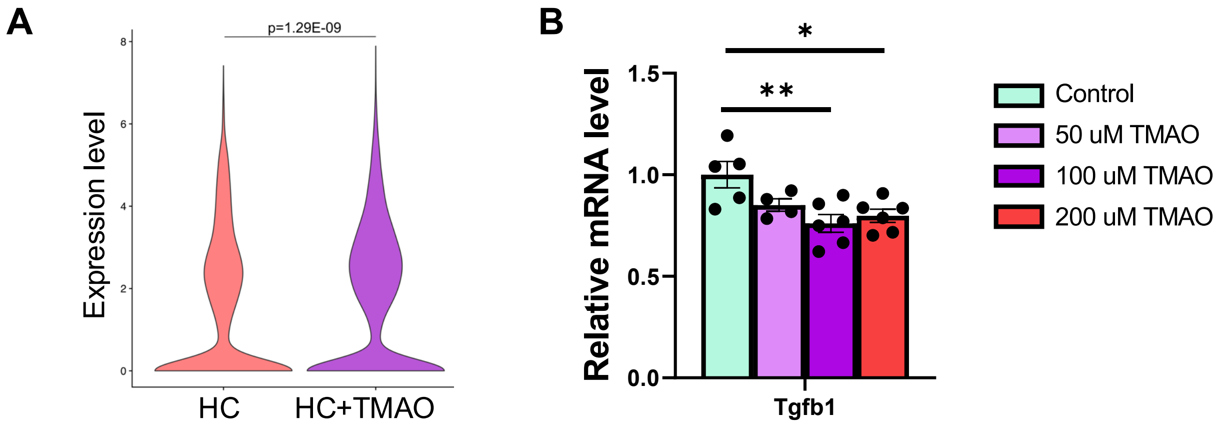

**Table S1. qPCR primers used in the study.**

| Gene Name | Target Species | Fwd Primer 5' -> 3' | Rev Primer 5' -> 3' |
|---|---|---|---|
| ***ATP8*** | Human | ATACTACCGTATGGCCCACC | GGGCTTTGGTGAGGGAGGTA |
| ***B2M*** | Human | AGATGAGTATGCCTGCCGTG | TCATCCAATCCAAATGCGGC |
| ***COL1A1*** | Human | GACATGTTCAGCTTTGTGGACC | TGGTCTCGTCACAGATCACG |
| ***COL1A2*** | Human | TGGTCTCGGTGGGAACTTTG | CACCCTGTGGTCCAACAACT |
| ***ND3*** | Human | GCGGCTTCGACCCTATATCC | AGGGCTCATGGTAGGGGTAA |
| ***SRSF4*** | Human | TGTGGAGCGCTTCTTTAAGGG | ATAACCATATCCACTGCGTCCA |
| ***TGFB1*** | Human | TACCTGAACCCGTGTTGCTC | CCGGTAGTGAACCCGTTGAT |
| ***TP53INP1*** | Human | CGCTGCACAACAACAAAAGGA | AGCAAGAAGAGTCATTGTACGTGG |
| ***Atp8*** | Mouse | CAAACATTCCCACTGGCACC | TTGTTGGGGTAATGAATGAGGC |
| ***Nd3*** | Mouse | ACAAGCTCTGCACGTCTACC | TGCTCATGGTAGTGGAAGTAGAA |

| | | | |
|---|---|---|---|
| ***Rpl13a*** | Mouse | CCCTCCACCCTATGACA<br>AGA | TTCTCCTCCAGAGTGGCTG<br>T |
| **Tgfb1** | Mouse | AGCTGCGCTTGCAGAGA<br>TTA | AGCCCTGTATTCCGTCTCC<br>T |

**Table S2. HC vs. Chow DEGs (adjusted p-value<0.05) shared across and unique to aortic cell types recovered in scRNAseq.**

**Table S3. HC+TMAO vs. HC DEGs (adjusted p-value<0.05) shared across and unique to aortic cell types recovered in scRNAseq.**

**Table S4. HC+TMAO vs. HC DEGs (adjusted p-value<0.05) shared across and unique to macrophage subtypes.**

**Table S5. HC vs. Chow upregulated DEG pathway enrichment (FDR<0.05) with Reactome.**

**Table S6. HC vs. Chow downregulated DEG pathway enrichment (FDR<0.05) with Reactome.**

**Table S7. TMAO+HC vs. HC upregulated DEG pathway enrichment (FDR<0.05) with Reactome.**

**Table S8. TMAO+HC vs. HC downregulated DEG pathway enrichment (FDR<0.05) with Reactome.**

**Table S9. TMAO+HC vs. HC DEG pathway enrichment (FDR<0.05) with Reactome. DEGs with p-value<0.01 were used.**
